## Supplemental figures and tables for "Immune correlates of early clearance of *Mycobacterium tuberculosis* among tuberculosis household contacts in Indonesia"

### Table S1. Clinical characteristics of IGRA-negative tuberculosis household contacts

| | Total | | | Subset[d] | | |
|---|---|---|---|---|---|---|
| | IGRA converters[a] (N = 116) | Persistently IGRA-negatives[a] (N = 317) | *P* value[b] | True IGRA converters[a] (N = 51) | True persistently IGRA-negatives[a] (N = 237) | *P* value[b] |
| **Case contact characteristics** | | | | | | |
| Age | 23 (15 – 36) | 22 (12 – 40) | 0.8 | 19 (11 – 32) | 21 (11 – 38) | 0.5 |
| Female sex | 51% | 54% | | 57% | 54% | |
| Presence of BCG scar | 74% | 87% | <0.001 | 69% | 90% | <0.001 |
| Current and previous smoking | 36% | 29% | 0.2 | 31%) | 25% | 0.4 |
| BMI, kg/$m^2$ | 21.0 (17.7 – 24.8) | 20.0 (16.7 – 24.3) | 0.12 | 19.8 (16.6 – 23.9) | 19.9 (16.5 – 24.2) | >0.9 |
| Diabetes[c] | 5.2% | 3.5% | 0.5 | 3.9% | 3.8% | >0.9 |
| **Exposure to the index case** | | | | | | |
| Exposure risk score[e] | 0.32 (0.23 – 0.38) | 0.25 (0.17 – 0.32) | <0.001 | 0.32 (0.25 – 0.38) | 0.23 (0.17 – 0.30) | <0.001 |
| Sleeping in the same room as the index case | 19% | 20% | 0.8 | 18% | 16% | 0.7 |
| Waking hours spent with the index case a day before enrollment | 5 (3 – 8) | 3 (1 – 7) | 0.003 | 4 (3 – 9) | 3 (1 – 6) | 0.026 |
| Index case highest smear grade | | | <0.001 | | | <0.001 |
| *Scanty* | 2.6% | 10% | | 0% | 10% | |
| *1+* | 21% | 31% | | 14% | 35% | |
| *2+* | 24% | 26% | | 27% | 25% | |
| *3+* | 53% | 33% | | 59% | 30% | |
| Presence of cavities on chest x-ray of index | 50% | 42% | 0.2 | 48% | 38% | 0.2 |
| Extent of x-ray abnormalities | 40 (25 – 55) | 35 (20 – 60) | 0.6 | 40 (25 – 64) | 35 (25 – 55) | 0.7 |
| *M. tuberculosis* Beijing genotype in the index case | 36% | 25% | 0.028 | 40% | 23% | 0.016 |
| Blood count parameters at baseline | | | | | | |
| Hemoglobin g/dL | 14.15 (13.10 – 15.43) | 13.60 (12.80 – 14.90) | 0.024 | 13.90 (12.95, 14.85) | 13.50 (12.80, 14.60) | 0.4 |
| Platelets 1,000/$mm^3$ | 298 (249 – 362) | 306 (261 – 359) | 0.8 | 305 (267 – 370) | 308 (266 – 362) | 0.7 |
| Leukocytes 1,000/$mm^3$ | 7.55 (6.20 – 8.63) | 7.40 (6.30 – 8.50) | >0.9 | 7.60 (6.40 – 8.50) | 7.30 (6.20 – 8.50) | 0.7 |
| Lymphocytes 1,000/µL | 2.56 (2.13 – 3.13) | 2.64 (2.08 – 3.06) | >0.9 | 2.72 (2.20 – 3.17) | 2.61 (2.02 – 3.02) | 0.2 |
| Neutrophiles 1,000/µL | 4.12 (3.20 – 5.04) | 3.91 (3.20 – 5.01) | 0.8 | 4.16 (3.28 – 4.85) | 3.90 (3.24 – 4.99) | >0.9 |
| Monocytes 1,000/µL | 0.42 (0.33 – 0.54) | 0.43 (0.31 – 0.57) | 0.6 | 0.42 (0.34 – 0.53) | 0.43 (0.32 – 0.57) | 0.6 |
| Quantitative IFNγ release assay result | | | | | | |
| IFNγ Nil tube IU/L | 0.15 (0.08 – 0.31) | 0.13 (0.08 – 0.25) | 0.3 | 0.14 (0.07, 0.32) | 0.13 (0.08 – 0.24) | 0.5 |
| IFNγ TB-Nil tube IU/L | 0.08 (-0.01 – 0.23) | 0.01 (-0.02 – 0.08) | <0.001 | 0.00 (-0.08, 0.04) | 0.00 (-0.04 – 0.03) | 0.7 |
| IFNγ Mitogen-Nil tube IU/L | 8.08 (3.34 – 10.00) | 8.91 (3.42 – 10.00) | 0.5 | 8.93 (4.20, 10.00) | 8.51 (3.05 – 10.00) | 0.7 |

Abbreviations: BCG, Bacillus Calmette-Guerin; BMI, body mass index; IQR, interquartile range.

[a] Median (IQR); %

[b] Mann-Whitney U test; Pearson's Chi-squared test; Fisher's exact test

[c] Diabetes defined as follows: no diabetes, random capillary blood glucose >101 mg/dL or hemoglobin A1c (HbA1c) <5.7%; prediabetes, HbA1c 5.7%–6.4%; diabetes, HbA1c ≥6.5.

[d] Subset using strict IGRA cutoff of <0.15 IU/mL as negative and >0.70 IU/mL as positive result instead of the cutoff provided in the kit (0.35 IU/mL)

[e] Exposure risk scores were derived from a logistic regression model of *Mtb* exposure variables (index case: sputum smear grade, cavities, extent of x-ray disease; contacts: hours spent with and sleeping proximity to the case) with IGRA results at 14 weeks as the dependent variable[1]

#### Table S2. Relative Risk of IGRA conversion in contacts exposed to different strain and the effect of BCG vaccination using different cut-off criteria

**A**

**Relative Risk of IGRA conversion in contacts with manufacturer cut-off by index case *Mtb* genotype**

| Genotype | Persistently IGRA-negatives n=275 | IGRA converters n=108 | Relative Risk (95% CI) | *P* value | Adjusted RR (95% CI) | *P* value |
|---|---|---|---|---|---|---|
| Other | 206 (75%) | 69 (63%) | 1.00 (ref) | | 1.00 (ref) | |
| Beijing | 69 (25%) | 39 (37%) | 1.44 (0.98-2.10) | <0.001 | 1.39 (1.00-1.93) | 0.048 |

**Relative Risk of IGRA conversion in contacts *with strict cut-off* by index case *Mtb* genotype**

| Genotype | Persistently IGRA-negatives n=206 | IGRA converters n=50 | Relative Risk (95% CI) | *P* value |
|---|---|---|---|---|
| Other | 158 (76%) | 30 (60%) | 1.00 (ref) | |
| Beijing | 48 (24%) | 20 (40%) | 1.84 (1.11-2.97) | 0.015 |

**B**

**Relative Risk of IGRA conversion by contacts BCG vaccination status and index case *Mtb* genotype**

| Genotype | BCG | Persistently IGRA-negatives n=275 | IGRA converters n=108 | Relative Risk (95% CI) | *P* value | Adjusted RR (95% CI) | *P* value |
|---|---|---|---|---|---|---|---|
| Other | No | 22 (10%) | 21 (30%) | 1.00 (ref) | | 1.00 (ref) | |
| | Yes | 184 (90%) | 48 (70%) | 0.42 (0.28-0.63) | <0.001 | 0.40 (0.27-0.61) | <0.001 |
| Beijing | No | 13 (19%) | 7 (18%) | 1.00 (ref) | | 1.00 (ref) | |
| | Yes | 56 (81%) | 32 (82%) | 1.04 (0.54-2.01) | 0.9 | 1.02 (0.56-1.85) | 0.9 |

**Relative Risk of IGRA conversion with strict IGRA cut-off by contacts BCG vaccination status and index case *Mtb* genotype**

| Genotype | BCG | Persistently IGRA-negatives n=206 | IGRA converters n=50 | Relative Risk (95% CI) | *P* value |
|---|---|---|---|---|---|
| Other | No | 14 (10%) | 12 (30%) | 1.00 (ref) | |
| | Yes | 144 (90%) | 18 (70%) | 0.24 (0.13-0.43) | <0.001 |
| Beijing | No | 6 (13%) | 4 (20%) | 1.00 (ref) | |
| | Yes | 42 (87%) | 16 (80%) | 0.69 (0.29-1.63) | 0.4 |

#### Table S3. Cytokine measurements of IGRA converters and persistently IGRA-negatives

| | IGRA converters (log10 pg/mL) N = 91[a] | Persistently IGRA-negative (log10 pg/mL) N = 237[a] | p-value[b] | q-value[c] |
|---|---|---|---|---|
| *E. coli* stimulation | | | | |
| TNF_E.coli | 2.56 (2.17 – 2.95) | 2.70 (2.36 – 3.01) | 0.020 | 0.12 |
| IL-8_E.coli | 3.43 (3.23 – 3.68) | 3.56 (3.33 – 3.77) | 0.009 | 0.077 |
| IL-6_E.coli | 3.55 (3.28 – 3.76) | 3.67 (3.47 – 3.87) | 0.002 | 0.042 |
| IL-1b_E.coli | 2.83 (2.42 – 3.04) | 2.88 (2.58 – 3.11) | 0.12 | 0.41 |
| IL-1Ra_E.coli | 3.39 (3.19 – 3.53) | 3.41 (3.20 – 3.63) | 0.14 | 0.41 |
| IL-10_E.coli | 1.81 (1.39 – 2.01) | 1.88 (1.59 – 2.06) | 0.11 | 0.41 |
| BCG stimulation | | | | |
| TNF_BCG | 2.43 (2.19 – 2.69) | 2.47 (2.21 – 2.73) | 0.53 | 0.80 |
| IL-8_BCG | 4.12 (3.91 – 4.32) | 4.18 (3.99 – 4.33) | 0.38 | 0.62 |
| IL-6_BCG | 3.58 (3.42 – 3.76) | 3.64 (3.48 – 3.80) | 0.17 | 0.41 |
| IL-1b_BCG | 2.50 (2.26 – 2.77) | 2.49 (2.30 – 2.73) | 0.88 | 0.95 |
| IL-1Ra_BCG | 3.20 (3.08 – 3.38) | 3.25 (3.10 – 3.43) | 0.18 | 0.41 |
| IL-10_BCG | 1.56 (1.38 – 1.80) | 1.59 (1.43 – 1.73) | 0.95 | 0.95 |
| *Mtb* stimulation | | | | |
| TNF_Mtb | 2.25 (1.91 – 2.59) | 2.27 (2.01 – 2.57) | 0.59 | 0.81 |
| IL-8_Mtb | 4.20 (3.89 – 4.39) | 4.14 (3.84 – 4.39) | 0.86 | 0.95 |
| IL-6_Mtb | 3.47 (3.21 – 3.67) | 3.50 (3.25 – 3.72) | 0.35 | 0.62 |
| IL-1b_Mtb | 2.41 (2.08 – 2.69) | 2.39 (2.05 – 2.70) | 0.67 | 0.86 |
| IL-1Ra_Mtb | 3.20 (3.04 – 3.36) | 3.21 (3.06 – 3.39) | 0.32 | 0.62 |
| IL-10_Mtb | 1.35 (1.10 – 1.70) | 1.39 (1.17 – 1.60) | 0.92 | 0.95 |

[a] Median (IQR)

[b] Wilcoxon rank sum test

[c] False discovery rate correction for multiple testing

#### Table S4. Characteristics of household contacts with anti-*Mtb* antibodies measured

| | **Baseline IGRA-positive**[a] (N = 100) | **Baseline IGRA-negative**[a] (N = 433) | ***P* value**[b] |
|---|---|---|---|
| **Case contact characteristics** | | | |
| Age | 30 (14 – 46) | 22 (12 – 39) | 0.069 |
| Female sex | 60% | 53% | 0.20 |
| Presence of BCG scar | 79% | 84% | 0.25 |
| Smoking | 32% | 31% | 0.21 |
| BMI, kg/m$^2$ | 20.8 (16.8 – 24.2) | 20.2 (16.8 – 24.4) | 0.72 |
| Diabetes[c] | 2.0% | 3.9% | 0.57 |
| Exposure to the index case | | | |
| Sleeping in the same room as the index case | 34% | 20% | 0.002 |
| Waking hours spent with the index case a day before enrollment | 6.0 (2.0 – 10.0) | 4.0 (1.0 – 8.0) | 0.006 |
| Index case highest smear grade | | | 0.17 |
| *Scanty* | 3.0% | 8.3% | |
| *1+* | 24% | 28% | |
| *2+* | 27% | 25% | |
| *3+* | 46% | 38% | |
| Presence of cavities on chest x-ray of index | 58% | 44% | 0.011 |
| Extent of x-ray abnormalities | 45 (25 – 66) | 40 (25 – 59) | 0.24 |
| *M. tuberculosis* Beijing genotype in the index case | 26% | 28% | 0.76 |
| Blood count parameters at baseline | | | |
| Hemoglobin g/dL | 13.55 (12.65 – 14.70) | 13.70 (12.80 – 15.00) | 0.17 |
| Platelets 1,000/mm$^3$ | 289 (249 – 339) | 305 (258 – 360) | 0.16 |
| Leukocytes 1,000/mm$^3$ | 7.45 (6.48 – 8.50) | 7.40 (6.20 – 8.60) | 0.91 |
| Lymphocytes 1,000/µL | 2.70 (2.15 – 3.26) | 2.60 (2.12 – 3.07) | 0.17 |
| Neutrophiles 1,000/µL | 3.98 (3.29 – 4.81) | 4.03 (3.20 – 5.02) | 0.81 |
| Monocytes 1,000/µL | 0.41 (0.30 – 0.55) | 0.43 (0.32 – 0.56) | 0.75 |
| Quantitative IFNγ release assay result at baseline | | | |
| IFNγ Nil tube IU/L | 0.15 (0.10 – 0.30) | 0.14 (0.08 – 0.28) | 0.24 |
| IFNγ TB-Nil tube IU/L | 2.22 (0.95 – 6.75) | 0.02 (-0.02 – 0.13) | <0.001 |
| IFNγ Mitogen-Nil tube IU/L | 10.00 (3.72 – 10.00) | 8.68 (3.41 – 10.00) | 0.31 |

Abbreviations: BCG, Bacillus Calmette-Guerin; BMI, body mass index; IQR, interquartile range.

[a] Median (IQR); %

[b] Mann-Whitney U test; Pearson's Chi-squared test; Fisher's exact test

[c] Diabetes defined as follows: no diabetes, random capillary blood glucose >101 mg/dL or hemoglobin A1c (HbA1c) <5.7%; prediabetes, HbA1c 5.7%–6.4%; diabetes, HbA1c ≥6.5.

**Table S5. Characteristics of BCG-vaccinated volunteers with antibody and PBMC stimulation measurements**

| Characteristic | N = 298[1] |
|---|---|
| Age | 23 (20 – 25) |
| Female | 56% |
| Body mass index | 22.15 (20.80 – 23.62) |

[1] Median (IQR); n (%)

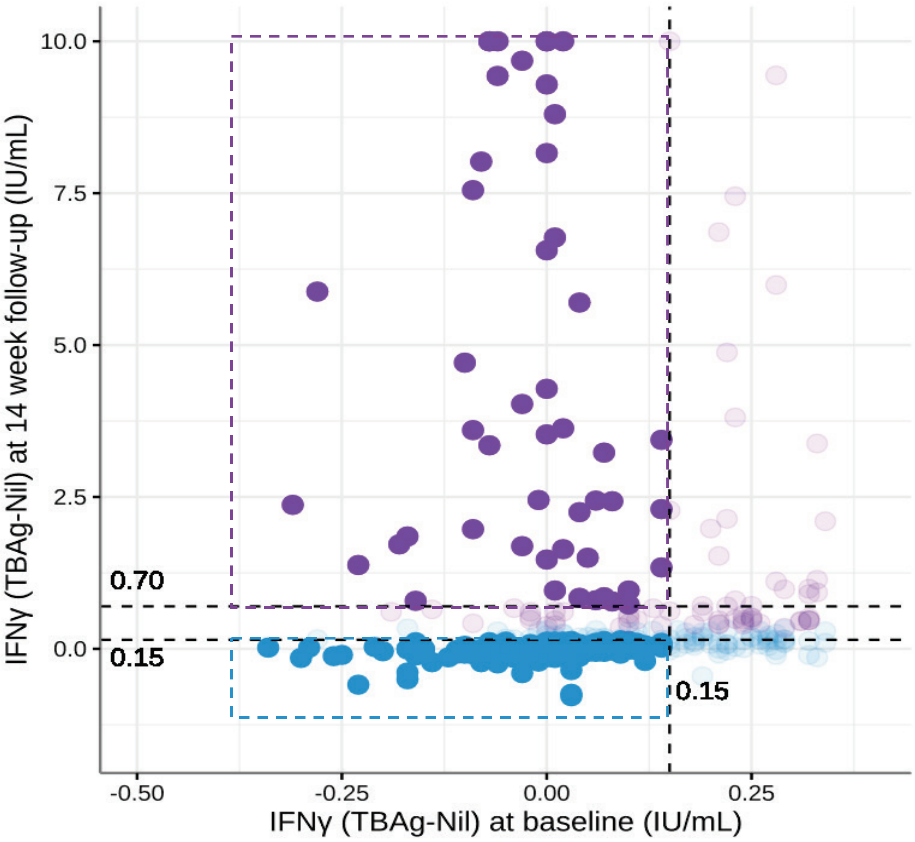


**Supplementary Figure 1. Subset of persistently IGRA-negative individuals and IGRA converters using stricter IGRA cutoff**

For IGRA negative individuals at baseline (N=433), we used a strict cut-off value for TB – Nil IFNγ of less than 0.15 IU/mL (both at baseline and 14 weeks), to classify subjects as persistently IGRA-negative (N=237, blue dotted box), and < 0.15 IU/mL at baseline and > 0.7 at 14 weeks to classify subjects as IGRA-converters (N=51, purple).

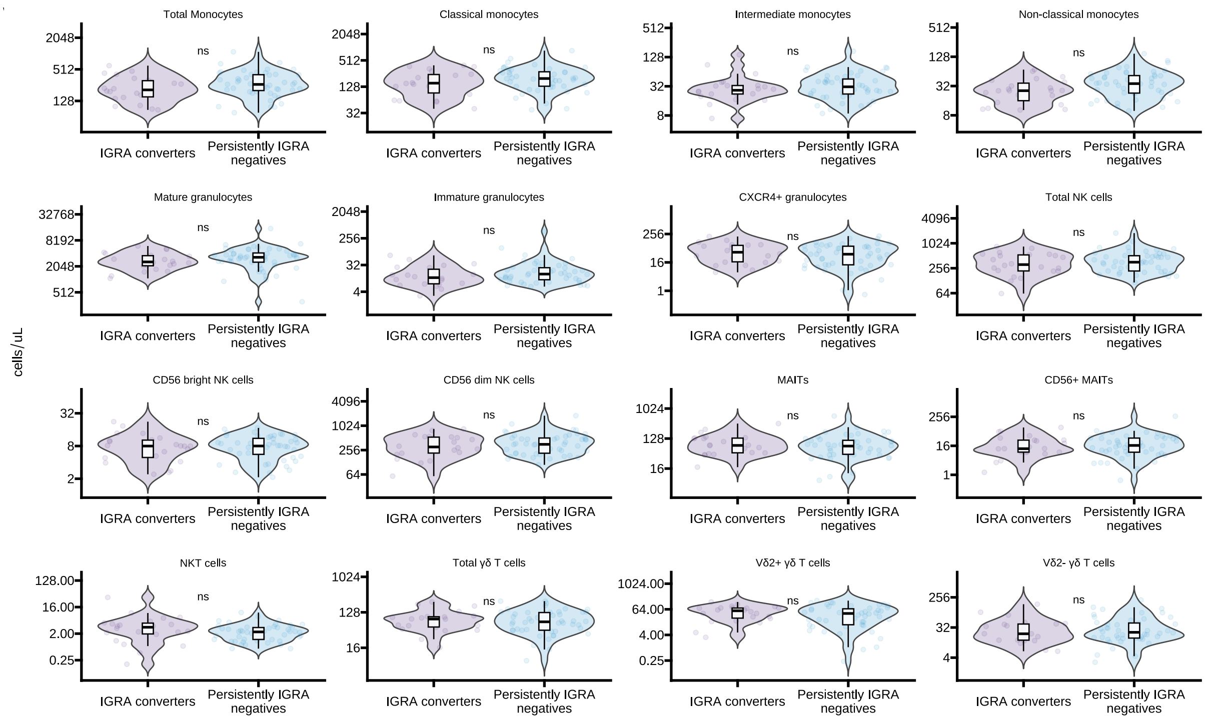


**Supplementary Figure 2. Innate immune cells in IGRA converters and persistently IGRA-negative individuals**

In IGRA negative individuals with complete flow-cytometric measurements, persistently IGRA-negative individuals (N = 48) and IGRA converters (N = 22), frequencies of circulating innate immune cells (numbers / µL blood) were compared at week 2. There was no difference between IGRA converters and persistently IGRA-negative individuals in individual innate immune cells (paired Mann-Whitney U test after Benjamini-Hochberg correction for multiple testing).

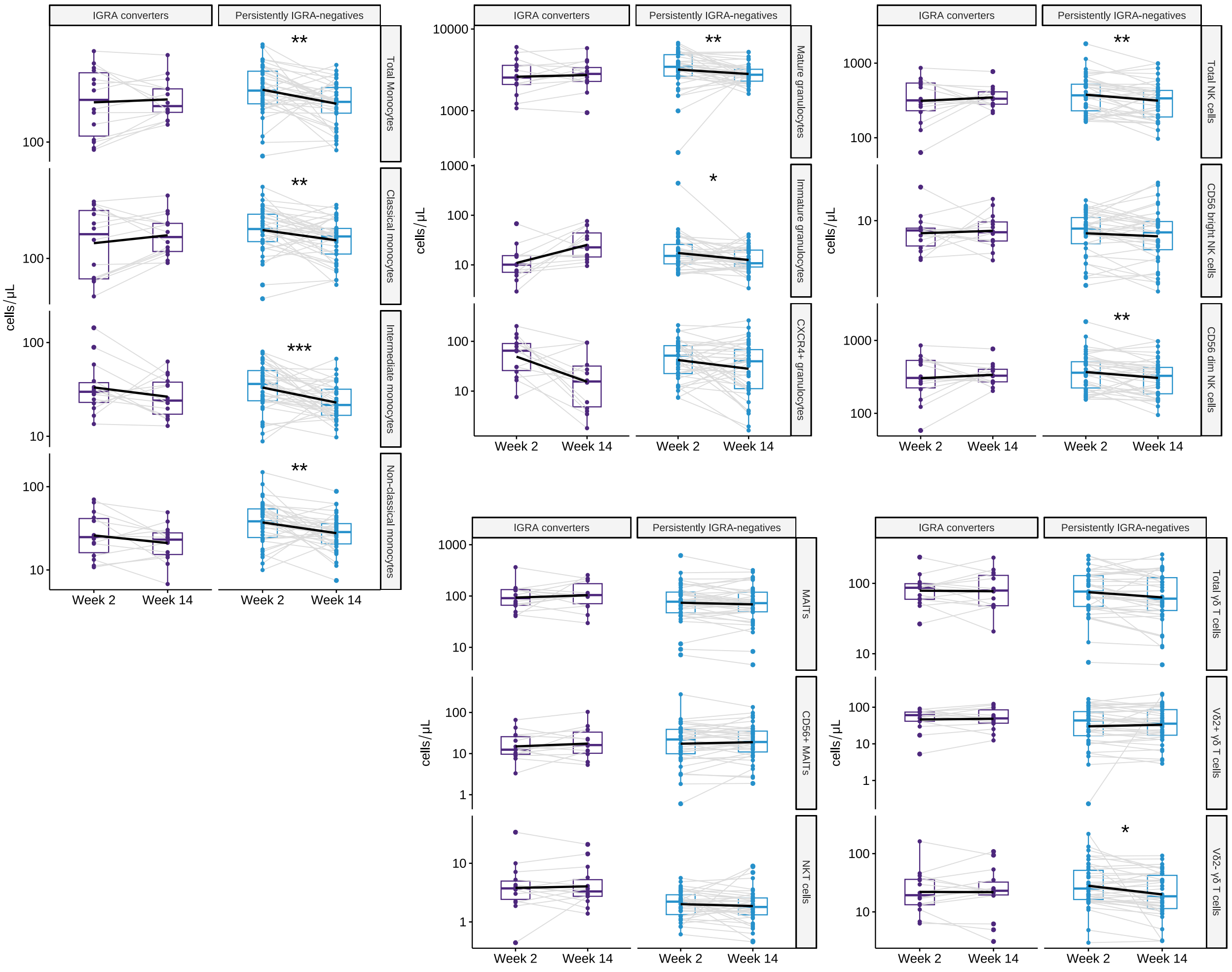


**Supplementary Figure 3. Innate immune cell population analysis of IGRA-negative individuals with BCG scar.**

In IGRA negative individuals with a BCG scar and complete flow-cytometric measurements, persistently IGRA-negative individuals (N = 38) and IGRA converters (N = 14), frequencies of circulating innate immune cells (numbers / μL blood) were compared between week 2 and week 14. In IGRA converters, no significant differences were seen (paired wilcoxon signed rank tests after Benjamini-Hochberg correction for multiple testing). In persistently IGRA-negative individuals, CD14$^{hi}$CD16$^{-}$ classical monocytes, CD14$^{hi}$CD16$^{+}$ intermediate monocytes, CD14$^{low}$CD16$^{+}$ non-classical monocytes, CD16$^{+}$ mature granulocytes, CD16$^{dim}$ immature granulocytes, and Vδ2$^{-}$ γδ T cells), were significantly lower at week 14 compared to week 2 and with a lower FDR compared to the complete dataset in Fig. 1, while CD56$^{dim}$ NK cells were also significantly lower. (FDR<0.1, <0.05, <0.01, <0.001; *, **, ***, ****)

**A**

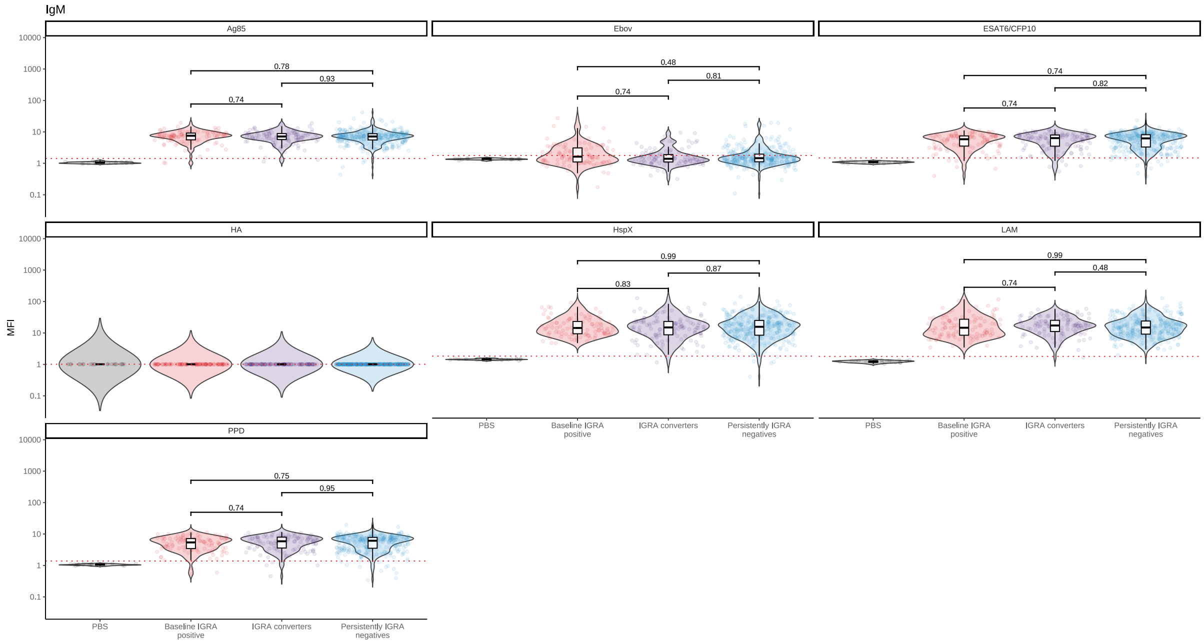


**Supplementary Figure 4. Antibody level as ratio to HA in all groups and PBS control**

Antibody levels ($log_{10}$ transformed), standardized to the positive control influenza virus hemaglutinnin (HA) as a ratio of MFI / MFI of anti-HA antibodies. To improve specificity (signal-noise ratio) the mean standardized antibody level + 6SD (standard deviation) in the PBS control was used as a cutoff (dotted lines), for IgM (**A**), total IgG (**B**), IgG1 (**C**), IgG2 (**D**), IgG3 (**E**), IgA1 (**F**), IgA2 (**G**), FcγR (**H-K**). Based on this cut-offs, we only included measurements of 5 antibody isotypes (IgM, IgG2, IgG3, IgA1, IgA2) against 5 different *Mtb* antigens in subsequent analysis. The antibody level as ratio to HA for Ebola-specific antibodies, as a negative control, were below the mean ratio to HA + 6 SD in PBS. All comparison were done using Mann-Whitney U test with correction for multiple testing using Benjamini-Hochberg (FDR).

**B**

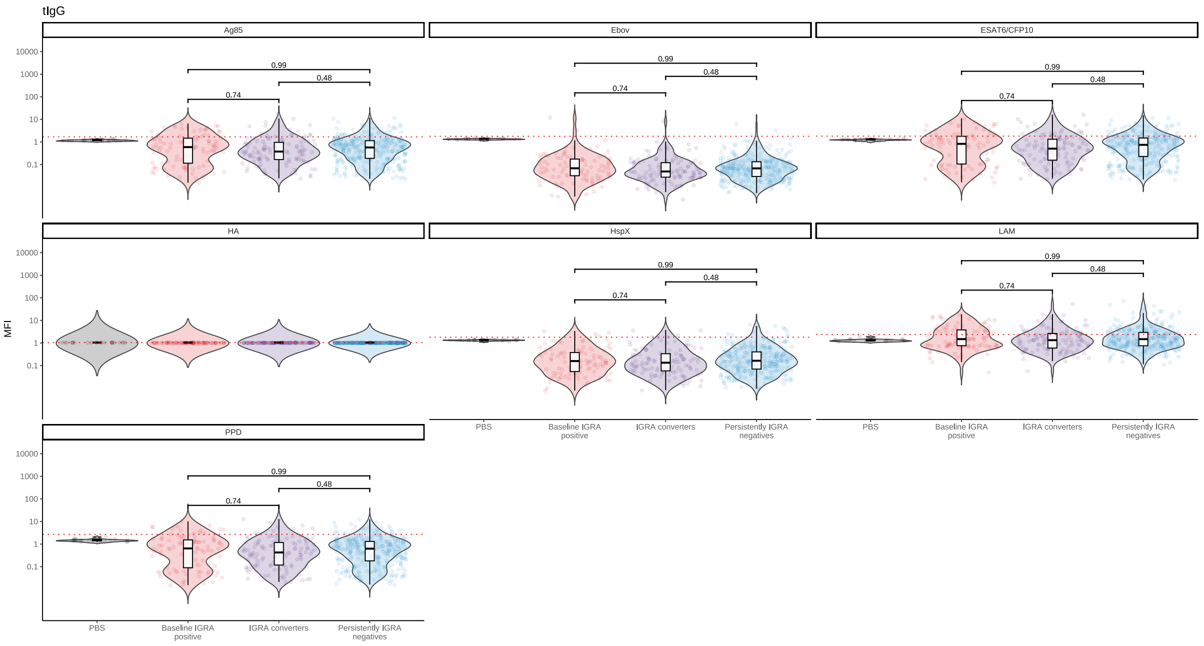
tIgG
Ag85
Ebov
ESAT6/CFP10
HA
HspX
LAM
PPD
MFI
10000
1000
100
10
1
0.1
0.99
0.48
0.74
PBS
Baseline IGRA positive
IGRA converters
Persistently IGRA negatives

**C**

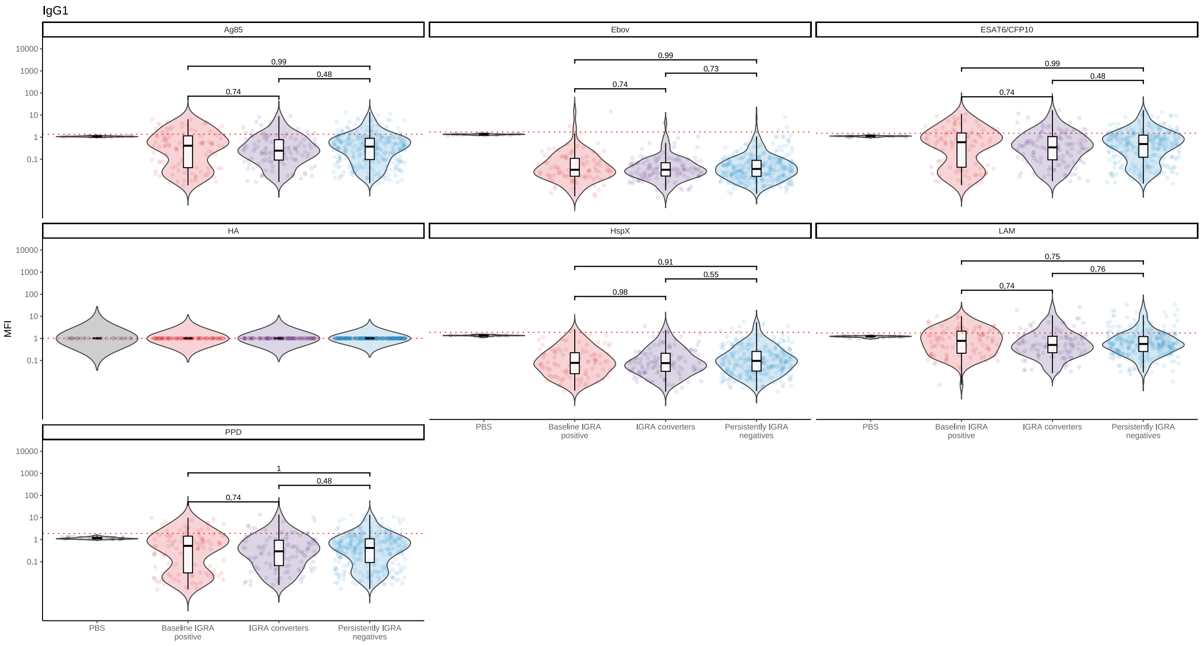
IgG1
Ag85
Ebov
ESAT6/CFP10
HA
HspX
LAM
PPD
MFI
10000
1000
100
10
1
0.1
0.99
0.48
0.74
0.73
0.91
0.55
0.98
0.75
0.76
PBS
Baseline IGRA positive
IGRA converters
Persistently IGRA negatives

D

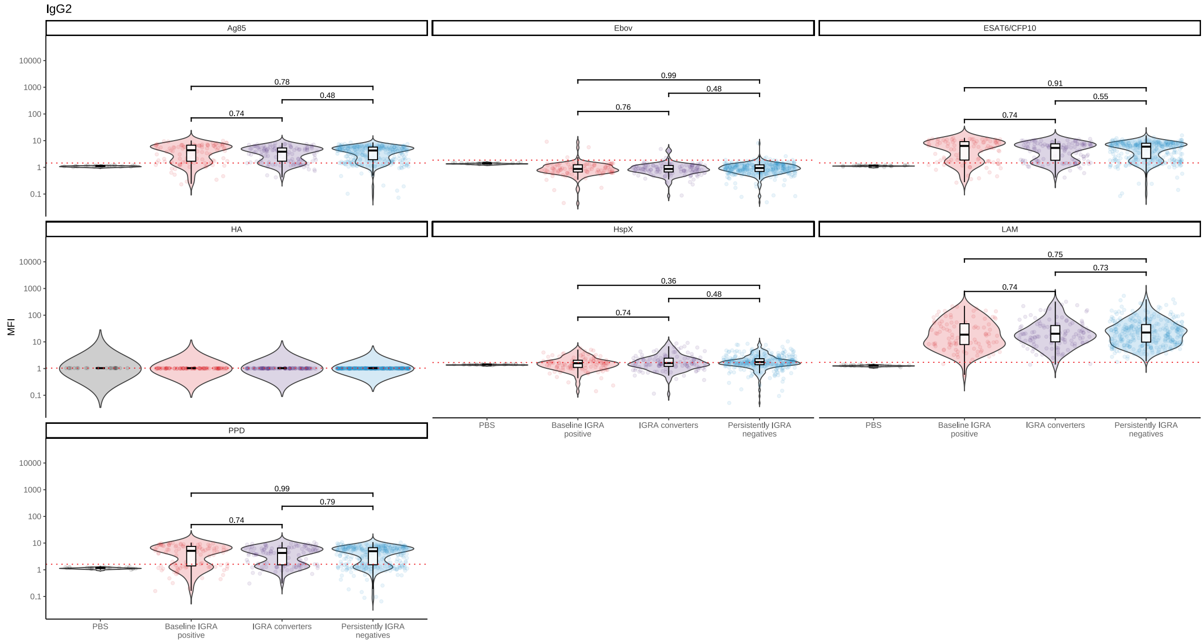
IgG2
Ag85
Ebov
ESAT6/CFP10
HA
HspX
LAM
PPD
MFI
10000
1000
100
10
1
0.1
0.78
0.48
0.74
0.99
0.48
0.76
0.91
0.55
0.74
0.36
0.48
0.74
0.75
0.73
0.74
0.99
0.79
0.74
PBS
Baseline IGRA positive
IGRA converters
Persistently IGRA negatives

E

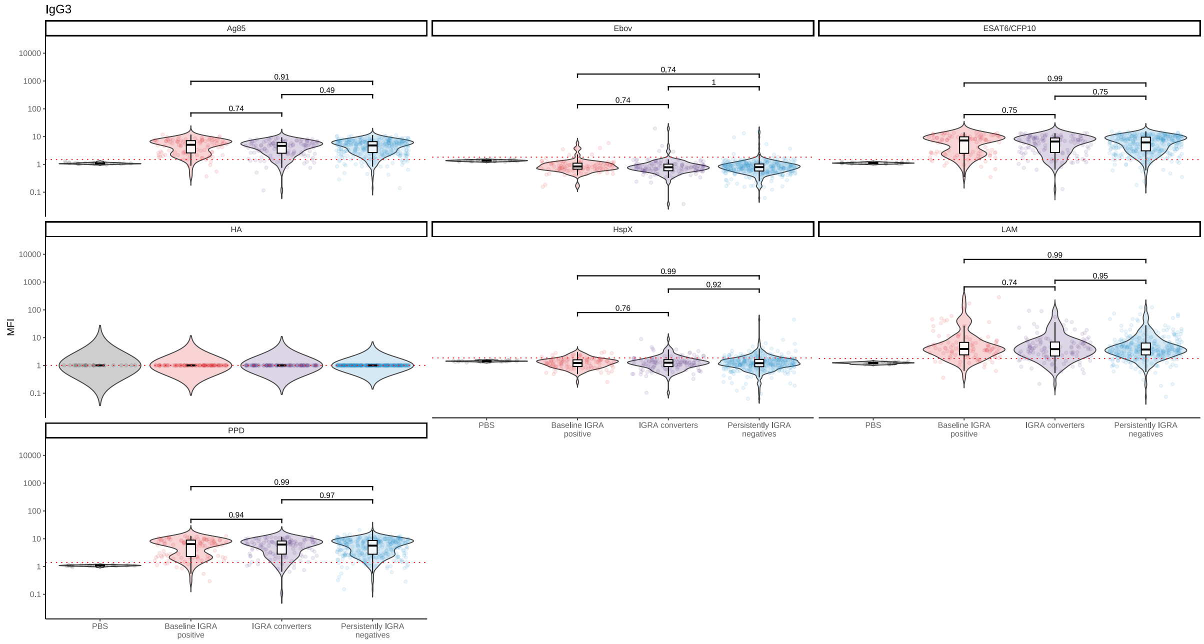
IgG3
Ag85
Ebov
ESAT6/CFP10
HA
HspX
LAM
PPD
MFI
10000
1000
100
10
1
0.1
0.91
0.49
0.74
0.74
1
0.74
0.99
0.75
0.75
0.99
0.92
0.76
0.99
0.74
0.95
0.99
0.97
0.94
PBS
Baseline IGRA positive
IGRA converters
Persistently IGRA negatives

F

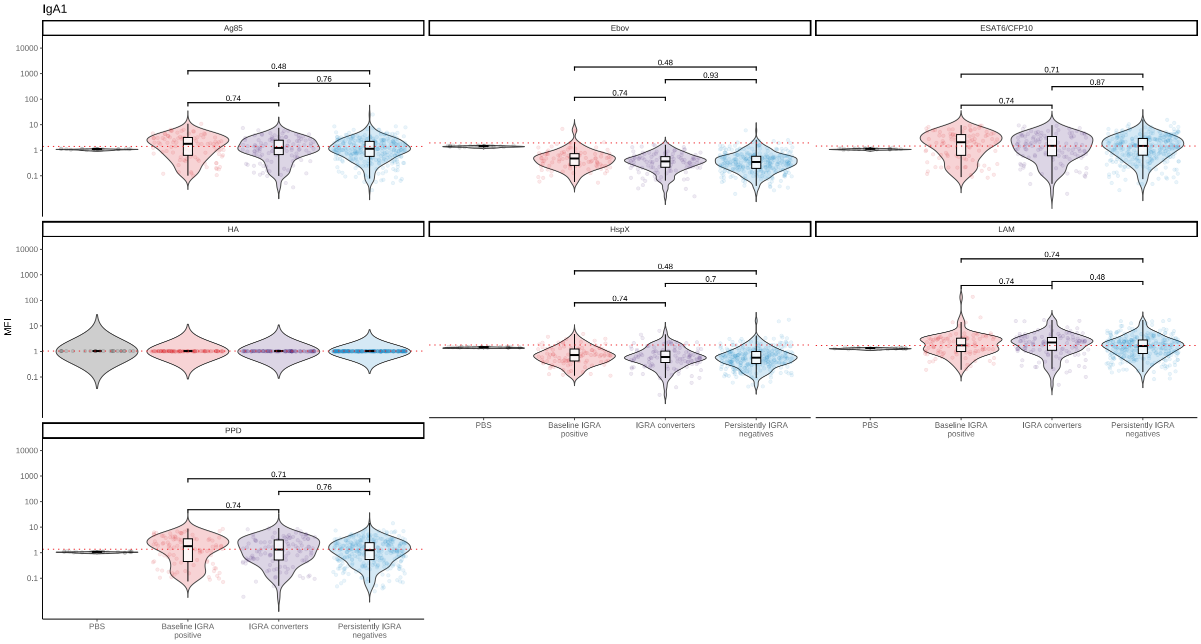
IgA1
Ag85
Ebov
ESAT6/CFP10
HA
HspX
LAM
PPD
MFI
10000
1000
100
10
1
0.1
0.48
0.76
0.74
0.93
0.71
0.87
0.7
0.48
PBS
Baseline IGRA positive
IGRA converters
Persistently IGRA negatives

G

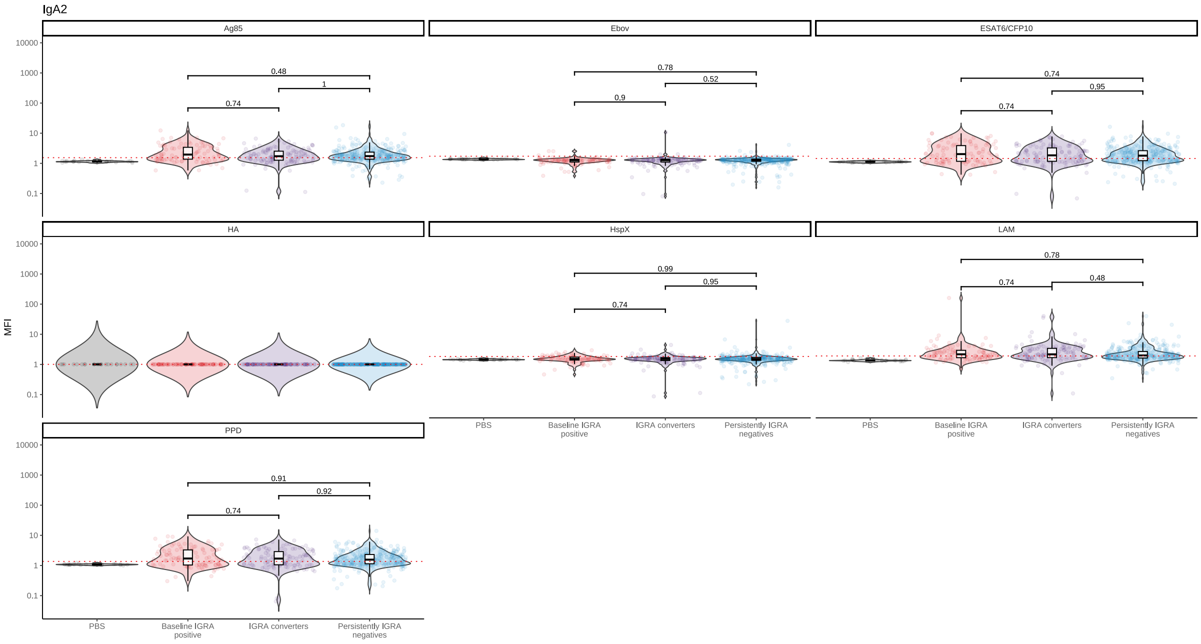
IgA2
Ag85
Ebov
ESAT6/CFP10
HA
HspX
LAM
PPD
MFI
10000
1000
100
10
1
0.1
0.48
1
0.74
0.78
0.52
0.9
0.95
0.99
0.48
0.91
0.92
PBS
Baseline IGRA positive
IGRA converters
Persistently IGRA negatives

H

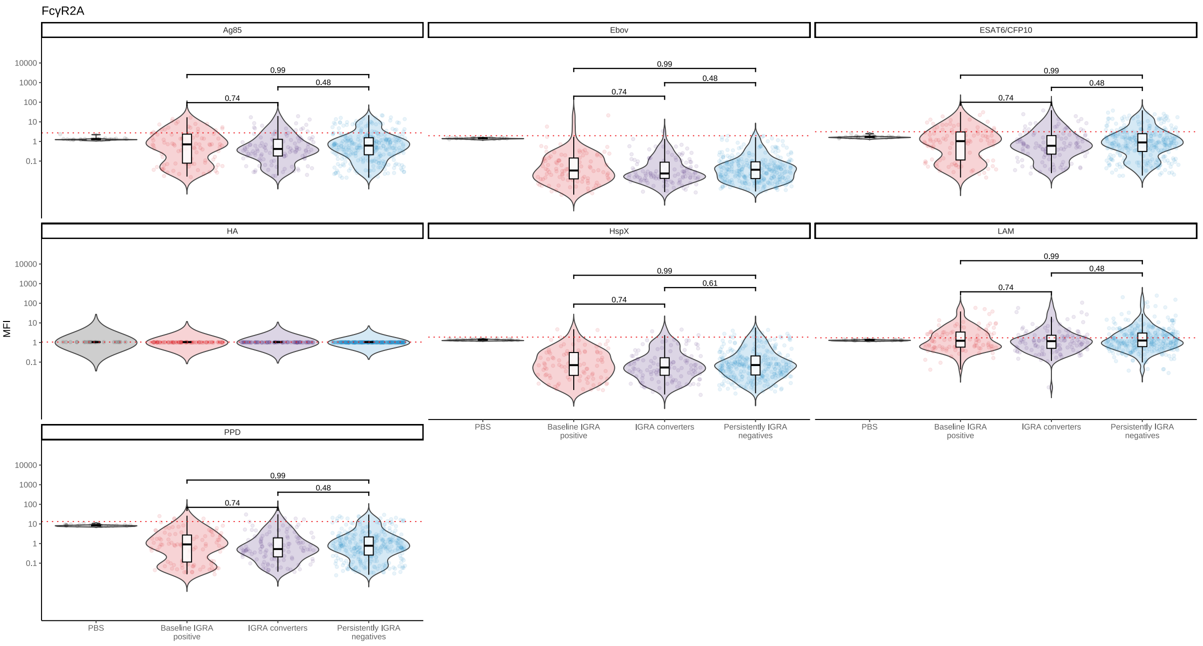
FcγR2A
Ag85
Ebov
ESAT6/CFP10
HA
HspX
LAM
PPD
MFI
PBS
Baseline IGRA positive
IGRA converters
Persistently IGRA negatives

I

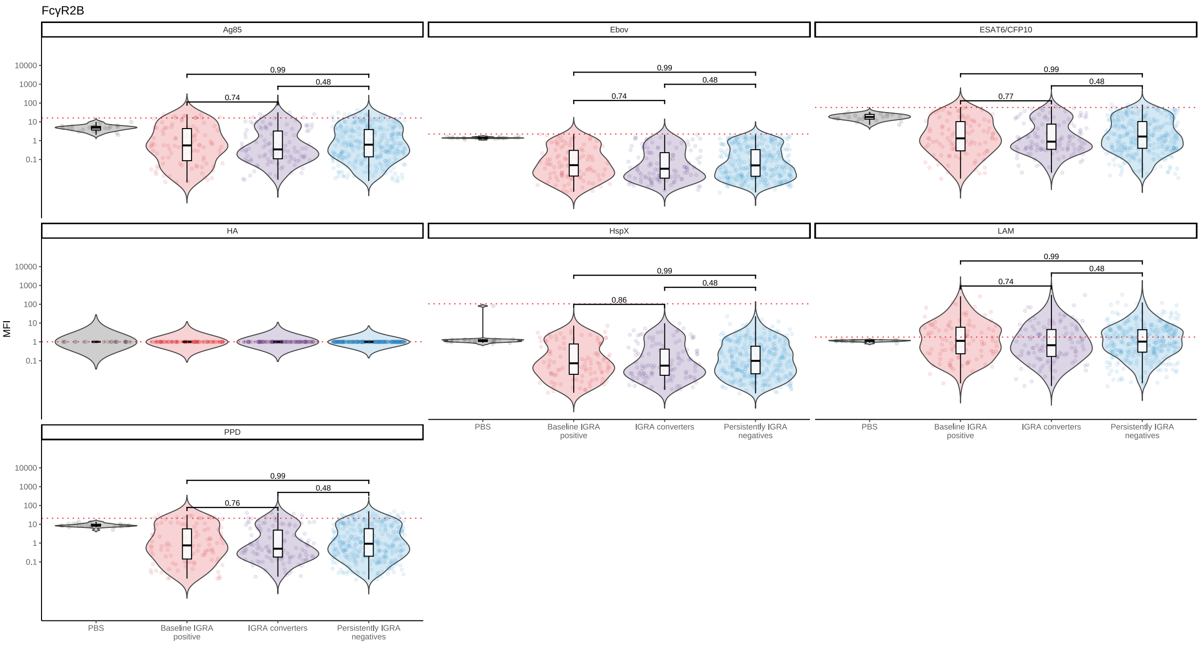
FcγR2B
Ag85
Ebov
ESAT6/CFP10
HA
HspX
LAM
PPD
MFI
PBS
Baseline IGRA positive
IGRA converters
Persistently IGRA negatives

**J**

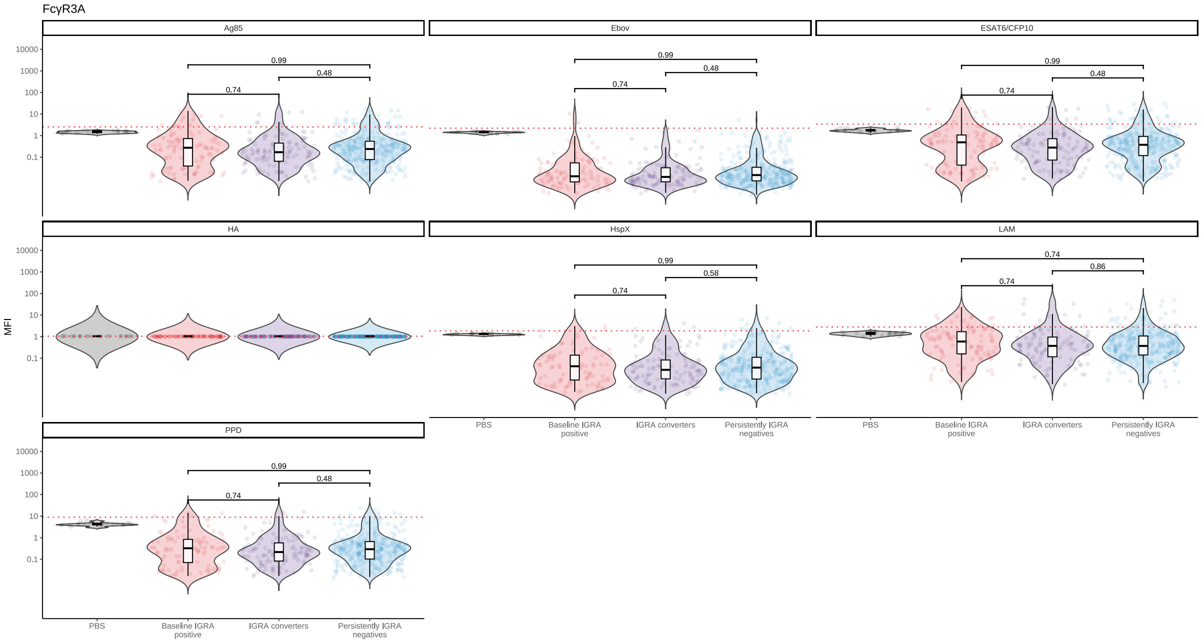
FcγR3A
Ag85
Ebov
ESAT6/CFP10
HA
HspX
LAM
PPD
MFI
10000
1000
100
10
1
0.1
0.99
0.48
0.74
0.58
0.86
PBS
Baseline IGRA positive
IGRA converters
Persistently IGRA negatives

**K**

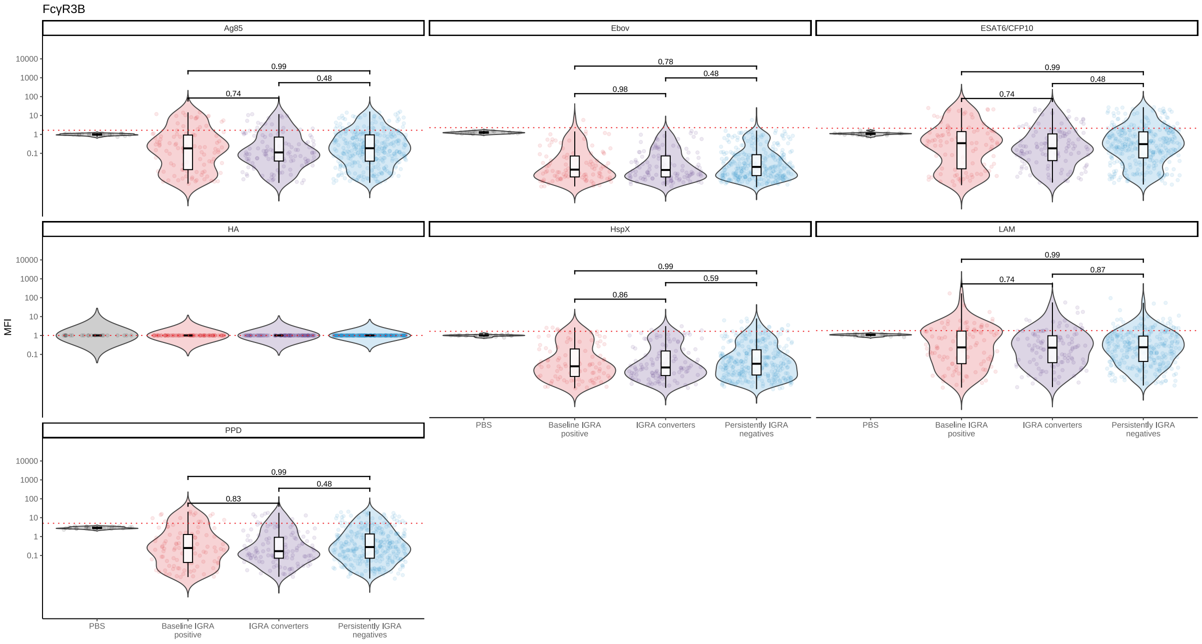
FcγR3B
Ag85
Ebov
ESAT6/CFP10
HA
HspX
LAM
PPD
MFI
10000
1000
100
10
1
0.1
0.99
0.48
0.74
0.78
0.98
0.86
0.59
0.87
0.83
PBS
Baseline IGRA positive
IGRA converters
Persistently IGRA negatives

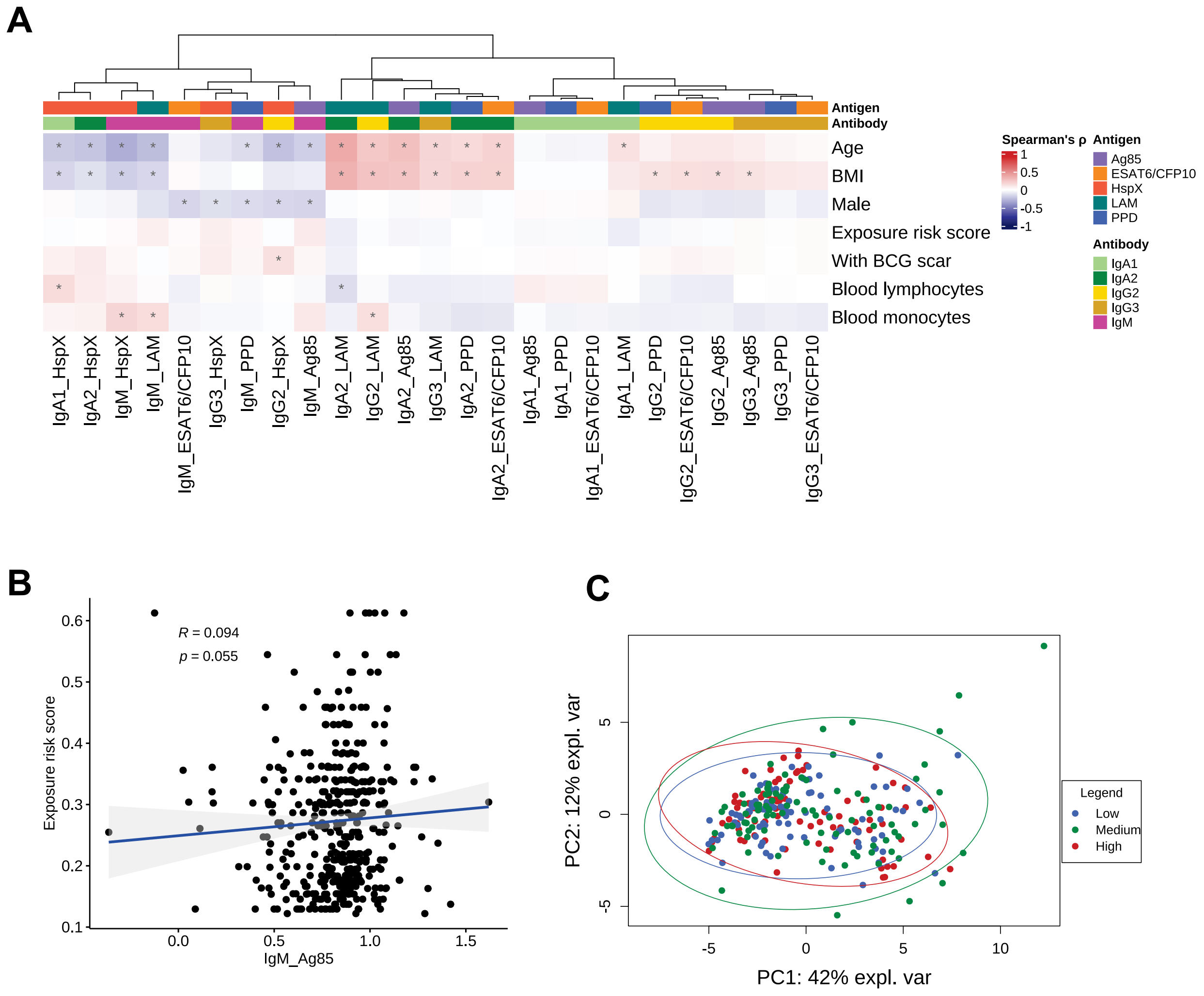


**Supplementary Figure 5. Correlation of antibody profiles with tuberculosis exposure and other subject characteristics in IGRA-negative individuals.**

Heatmap of the correlation between baseline antibody levels and subject characteristics, based on Spearman's rho (FDR <0.1, *) in IGRA-negative individuals (**A**). Correlation plot of Ag85-specific IgM and exposure risk score (**B**)**.** PCA of antibody measurements stratified by the level of exposure (Low = 1$^{st}$ tertile, Medium = 2$^{nd}$ tertile, High = 3$^{rd}$ tertile) (**C**).

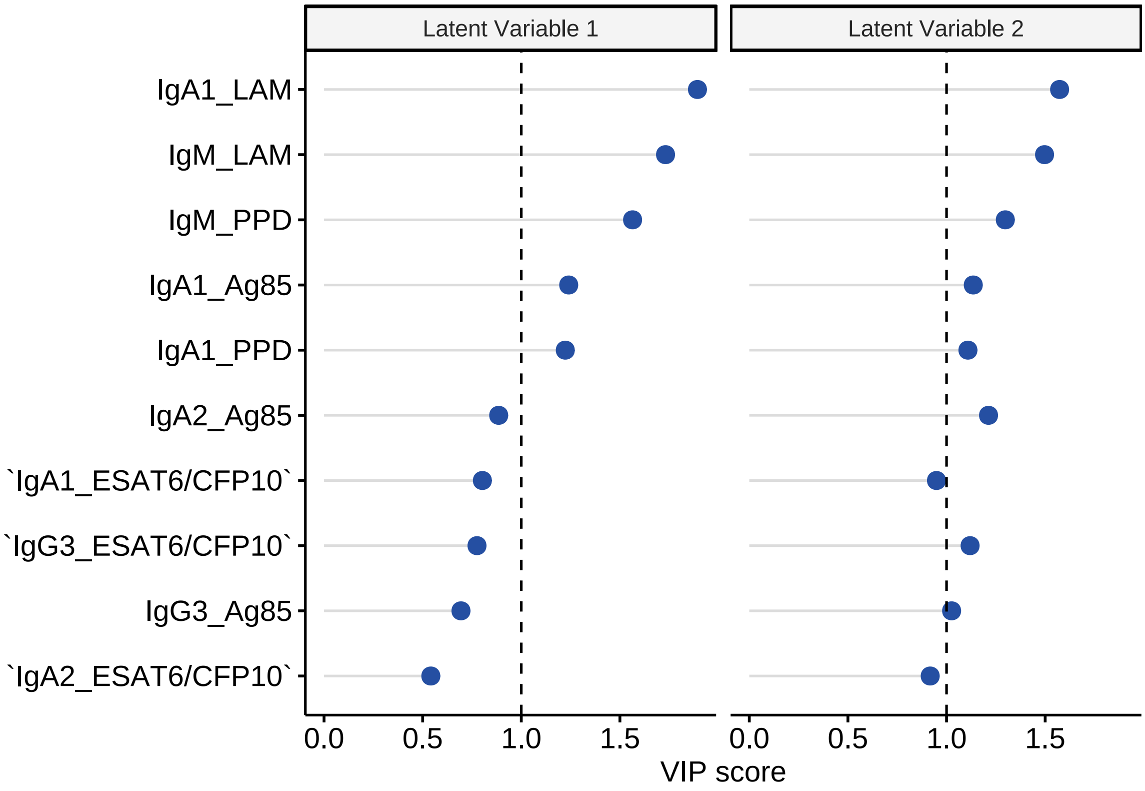


**Supplementary Figure 6. Variable importance in projection score of PLS-DA in IGRA-negative individuals.**

Variable importance in projection (VIP) coefficients of the PLS-DA in IGRA-negative individuals. The VIP score represent the relative importance of each antibody to explain IGRA converters and persistently IGRA-negatives group. The most relevant features to explain the difference between groups have VIP score > 1.

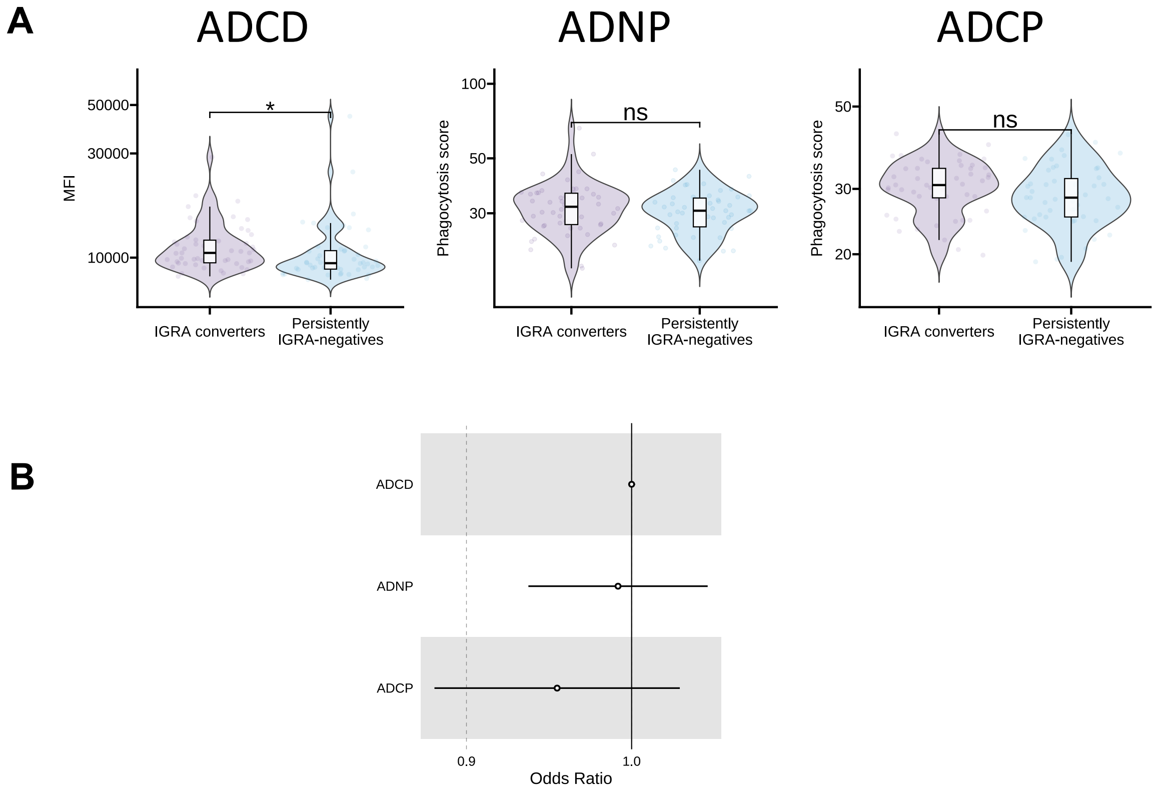


**Supplemental Figure 7. Antibody functionality in IGRA-converters and persistently IGRA-negative individuals.**

Using strict IGRA cutoff criteria (<0.15 IU/mL and >0.70 IU/mL), antibody dependent complement deposition, antibody-dependent cellular phagocytosis, and antibody-dependent neutrophil phagocytosis were compared between persistently IGRA-negatives (N = 50) and IGRA converters (N = 50). In Mann-Whitney U test, IGRA converters showed more antibody dependent complement deposition compared to persistently IGRA-negatives (FDR=0.031) (**A**), but in a logistic regression model adjusting for age, sex, BMI and correcting for multiple testing, there was no significant association between antibody function and IGRA conversion (open circle, FDR > 0.1) (**B**).

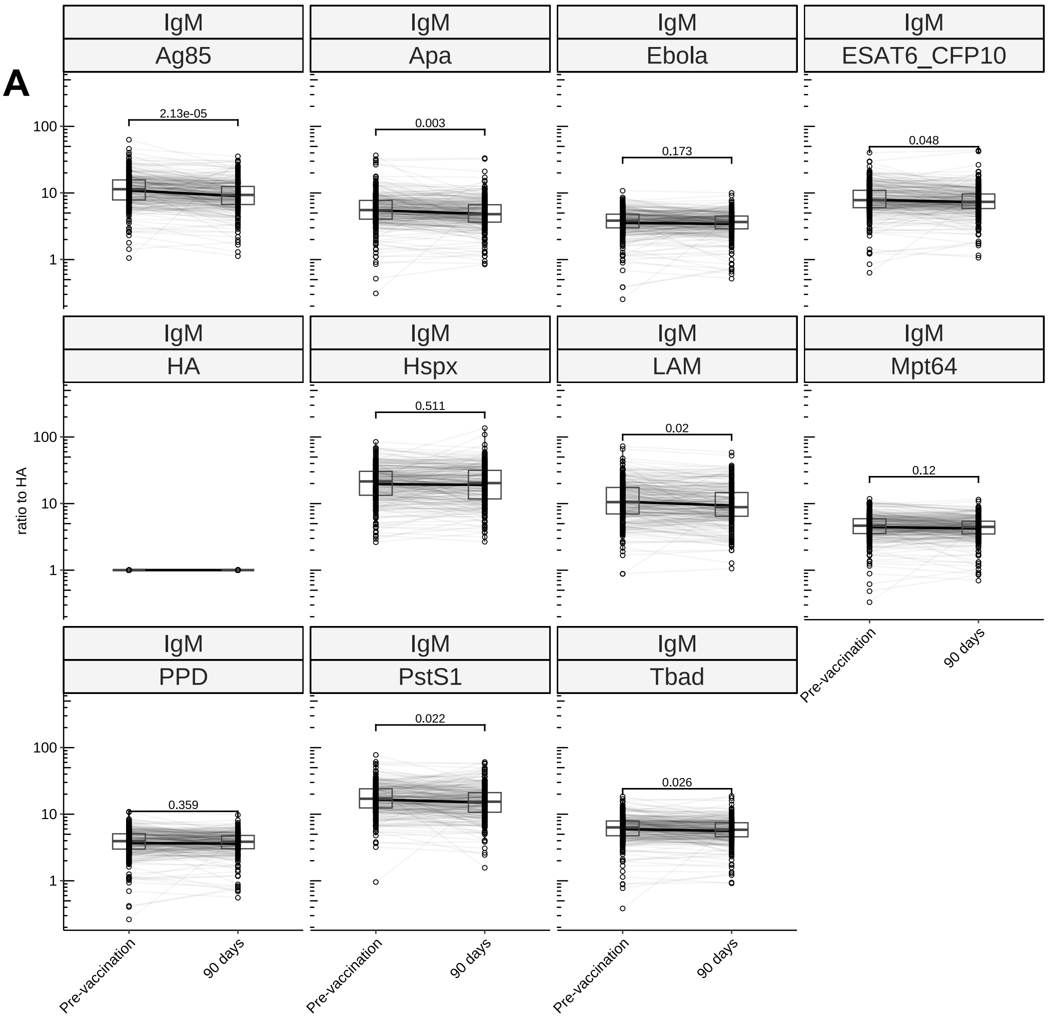


**Supplemental Figure 8. Effect of BCG vaccination on *Mtb* antigen-specific antibodies**

Concentrations of IgM (**A**), total IgG, IgG1 (**B**), IgG2, IgG3 (**C**), FcγR2A, FcγR3A (**D**) at baseline and 90 days after vaccination. (FDR<0.1, <0.05, <0.01, <0.001; *, **, ***, ****)

B

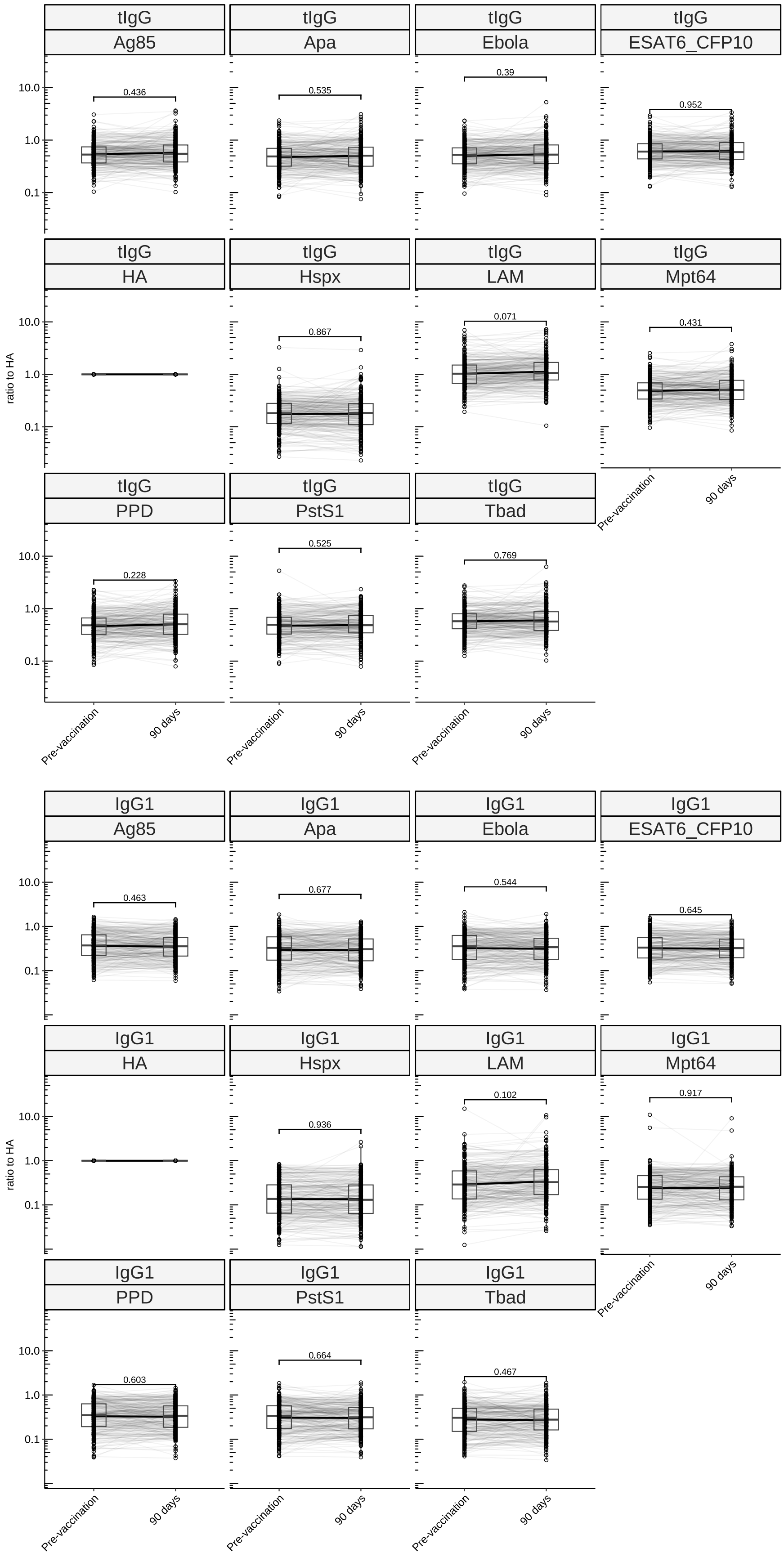
tIgG
Ag85
0.436
tIgG
Apa
0.535
tIgG
Ebola
0.39
tIgG
ESAT6_CFP10
0.952
tIgG
HA
tIgG
Hspx
0.867
tIgG
LAM
0.071
tIgG
Mpt64
0.431
tIgG
PPD
0.228
tIgG
PstS1
0.525
tIgG
Tbad
0.769
10.0
1.0
0.1
ratio to HA
Pre-vaccination
90 days
IgG1
Ag85
0.463
IgG1
Apa
0.677
IgG1
Ebola
0.544
IgG1
ESAT6_CFP10
0.645
IgG1
HA
IgG1
Hspx
0.936
IgG1
LAM
0.102
IgG1
Mpt64
0.917
IgG1
PPD
0.603
IgG1
PstS1
0.664
IgG1
Tbad
0.467

**C**

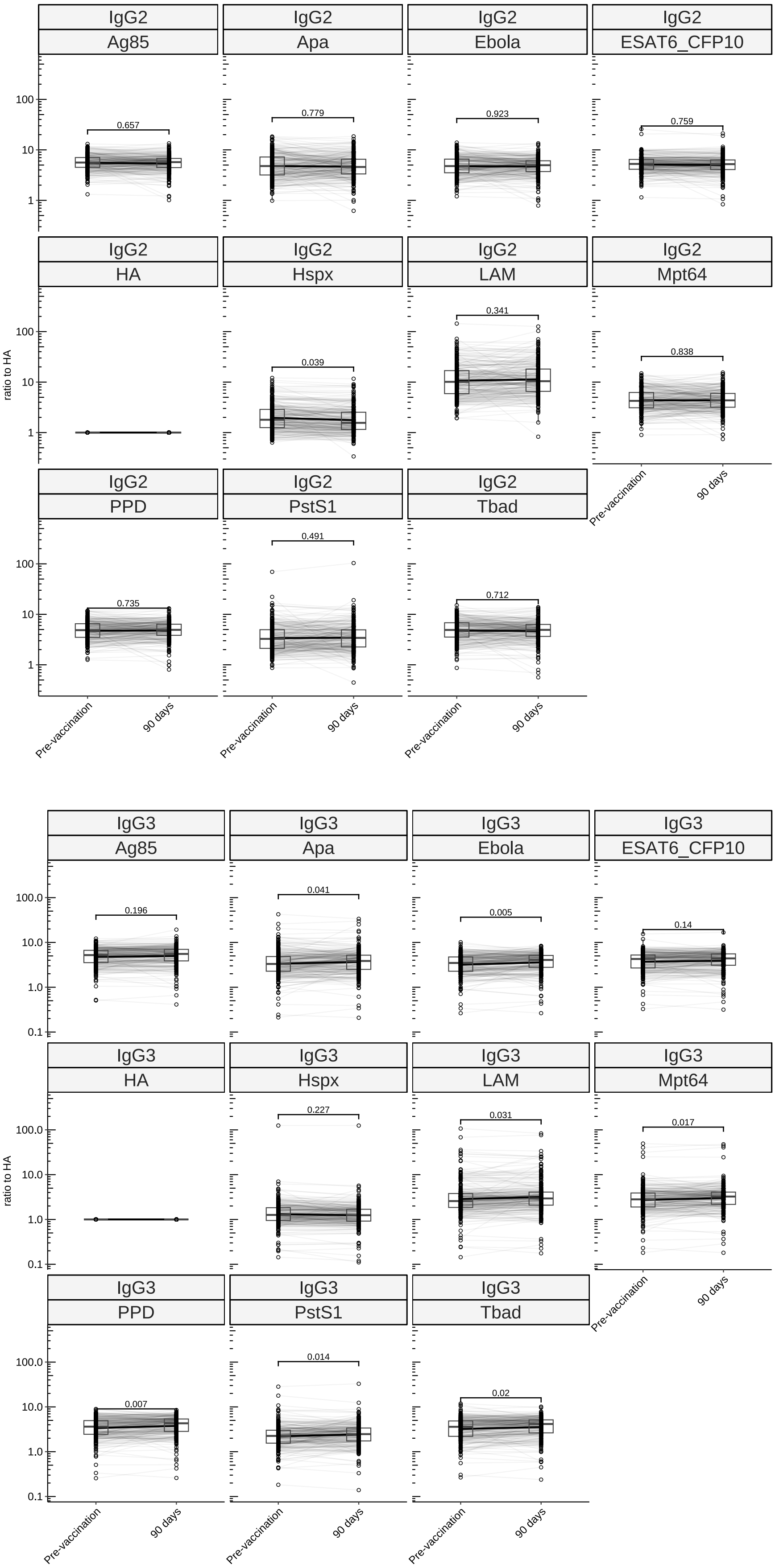

IgG2
Ag85
0.657
IgG2
Apa
0.779
IgG2
Ebola
0.923
IgG2
ESAT6_CFP10
0.759
IgG2
HA
IgG2
Hspx
0.039
IgG2
LAM
0.341
IgG2
Mpt64
0.838
IgG2
PPD
0.735
IgG2
PstS1
0.491
IgG2
Tbad
0.712
ratio to HA
100
10
1
Pre-vaccination
90 days
IgG3
Ag85
0.196
IgG3
Apa
0.041
IgG3
Ebola
0.005
IgG3
ESAT6_CFP10
0.14
IgG3
HA
IgG3
Hspx
0.227
IgG3
LAM
0.031
IgG3
Mpt64
0.017
IgG3
PPD
0.007
IgG3
PstS1
0.014
IgG3
Tbad
0.02
ratio to HA
100.0
10.0
1.0
0.1
Pre-vaccination
90 days

D

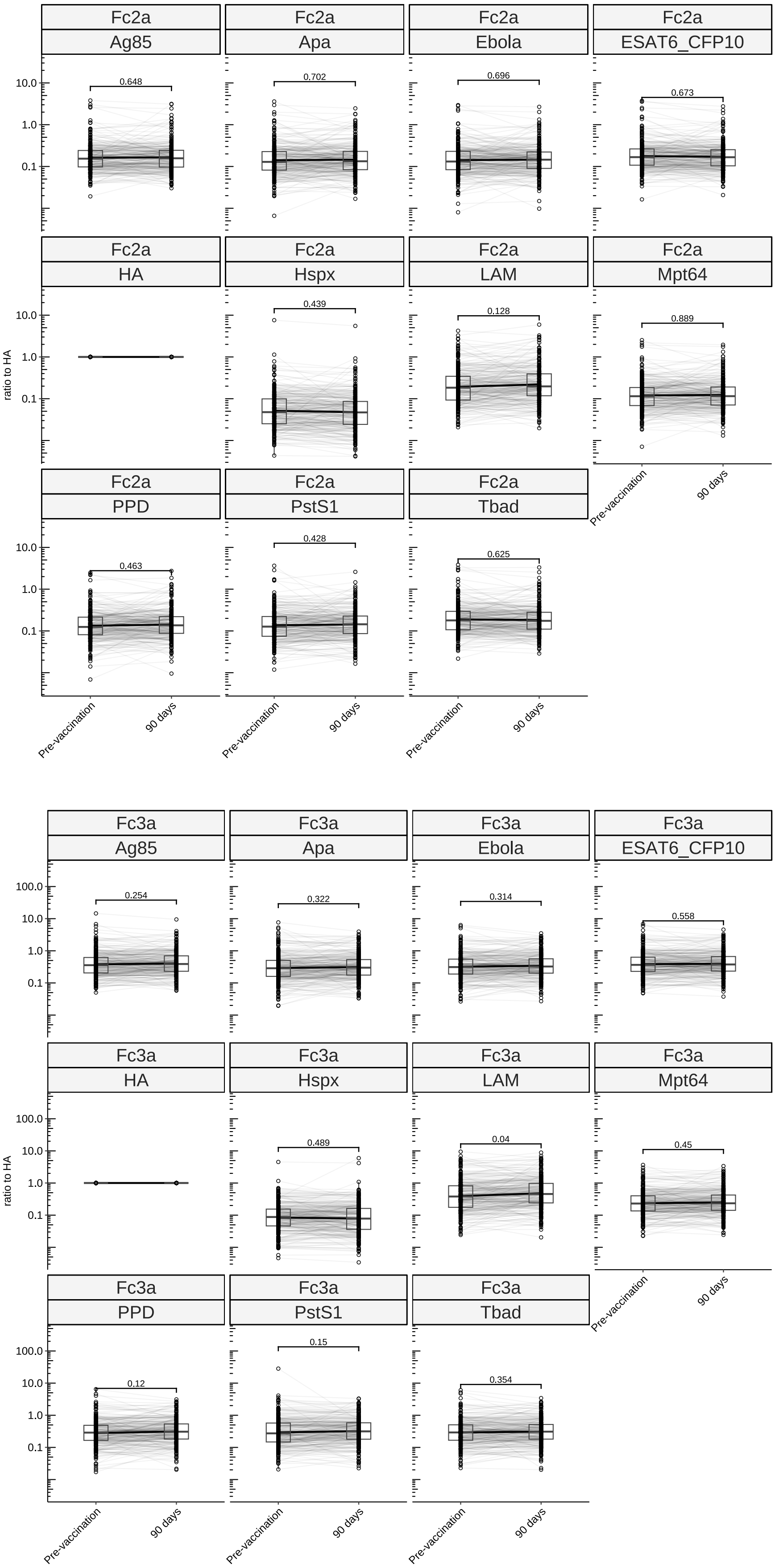
Fc2a
Ag85
0.648
Fc2a
Apa
0.702
Fc2a
Ebola
0.696
Fc2a
ESAT6_CFP10
0.673
Fc2a
HA
Fc2a
Hspx
0.439
Fc2a
LAM
0.128
Fc2a
Mpt64
0.889
Fc2a
PPD
0.463
Fc2a
PstS1
0.428
Fc2a
Tbad
0.625
10.0
1.0
0.1
ratio to HA
Pre-vaccination
90 days
Fc3a
Ag85
0.254
Fc3a
Apa
0.322
Fc3a
Ebola
0.314
Fc3a
ESAT6_CFP10
0.558
Fc3a
HA
Fc3a
Hspx
0.489
Fc3a
LAM
0.04
Fc3a
Mpt64
0.45
Fc3a
PPD
0.12
Fc3a
PstS1
0.15
Fc3a
Tbad
0.354
100.0
10.0
1.0
0.1
ratio to HA
Pre-vaccination
90 days
